# Selective inhibition of PfATP6 by artemisinins and identification of new classes of inhibitors after expression in yeast

**DOI:** 10.1101/2021.10.27.466210

**Authors:** Catherine M. Moore, Jigang Wang, Qingsong Lin, Pedro Ferreira, Mitchell A. Avery, Khaled Elokely, Henry M. Staines, Sanjeev Krishna

## Abstract

Treatment failures with artemisinin combination therapies (ACTs) threaten global efforts to eradicate malaria. They highlight the importance of identifying drug targets and new inhibitors and of studying how existing antimalarial classes work.

Herein we report the successful development of an heterologous expression-based compound screening tool. Validated drug target *P. falciparum* calcium ATPase6 (PfATP6) and a mammalian ortholog (SERCA1a) were functionally expressed in yeast providing a robust, sensitive, and specific screening tool. Whole-cell and *in vitro* assays consistently demonstrated inhibition and labelling of PfATP6 by artemisinins. Mutations in PfATP6 resulted in fitness costs that were ameliorated in the presence of artemisinin derivatives when studied in the yeast model.

As previously hypothesised, PfATP6 is a target of artemisinins. Mammalian SERCA1a can be mutated to become more susceptible to artemisinins. The inexpensive, low technology yeast screening platform has identified unrelated classes of druggable PfATP6 inhibitors. Resistance to artemisinins may depend on mechanisms that can concomitantly address multi-targeting by artemisinins and fitness costs of mutations that reduce artemisinin susceptibility.

## Introduction

The rate of decline in global cases of malaria has diminished in recent years^1^. Fortunately, artemisinin-containing antimalarial combination therapies (ACTs) continue to be effective in managing uncomplicated malaria, including in those regions with multidrug resistant parasites^2^. Artemisinins in ACTs remain effective even when their partner drug is failing^3^. In the past few years decreased parasite clearance times following treatment with ACTs have been associated with decreased sensitivity of *Plasmodium falciparum* ring stages to dihydroartemisinin (DHA)^4^. Prolonged parasite clearance times in and of themselves are not associated with treatment failures if the partner drug of an ACT is effective, but concerns have been raised about the risk of artemisinin resistance^5,6^.

Understanding how artemisinins act as antimalarials can help to optimise their use, increase insights into potential artemisinin resistance mechanisms, and help to develop them for other urgently needed indications such as their repurposing as anti-cancer^7^ or anti-SARS-CoV-2 agents^8^. Almost two decades ago, we suggested that artemisinins acted by inhibiting PfATP6, the SERCA pump of malarial parasites^9^. Several independent lines of evidence were consistent with this hypothesis, including the following observations. There are structural similarities between thapsigargin, a specific mammalian SERCA pump inhibitor, and artemisinins. Synthesis of thaperoxide that incorporates an endoperoxide bridge into thapsigargin confirmed that these structures were relatable, because antimalarial potency and inhibition of PfATP6 were simultaneously increased^10^. There was a positive correlation between inhibitor profiles for antimalarial action *in vitro* (whole cell assays) and after heterologous expression of PfATP6^9^. Artemisinin sensitivity in parasites transfected with PfATP6 mutant ^L263E^PfATP6 showed increased variability in sensitivity assays, with decreased sensitivity to artemisinin and dihydroartemisinin in *ex vivo* and *in vivo* experiments^11^. Fluorescent derivatives of artemisinin and thapsigargin co-localised in parasites^12^. PfATP6 is an essential gene in parasites as attempts to knock it out are lethal^13^. Studies in unrelated systems, such as in mammalian cancer cell lines have also demonstrated inhibition of SERCA by artemisinins^14,15^.

Heterologous expression of PfATP6 using *Xenopus* oocytes showed selective inhibition by artemisinins^9^. Others attempted to reproduce inhibition by artemisinins using purified and reconstituted PfATP6 in membrane vesicles and after expression in oocytes, but were unsuccessful^16^.

Our understanding of mechanisms of action of artemisinins has recently progressed through use of ‘click’ chemistry approaches. Artemisinins are alkylating agents that form covalent bonds (after activation) with their targets^17^. This has allowed the labelling of dozens of proteins in asexual stage *P. falciparum* and validation of some of them as potential targets of artemisinins. Two independent groups almost simultaneously identified several *P. falciparum* transporters by labelling with derivatised artemisinins. PfATP6 was included in this list of labelled proteins in both *in vivo* experiments^18,19^.

To advance these studies, we developed a yeast functional rescue assay using a codon optimised construct of PfATP6. This robust assay allows screening and study of PfATP6 inhibitors and comparisons with mammalian SERCA, mutational analyses, biochemical studies and scaling for higher throughput investigations. We have also translated findings from heterologous expression in yeast to studies in parasites to confirm their relevance and report findings using these approaches.

## Results

### Screening tool development and optimisation

*Saccharomyces cerevisiae* (K667) is hypersensitive to extracellular calcium as endogenous P-type calcium ATPases are inactive (PMC1) or deleted (PMR1). K667 is functionally rescued with heterologous P-type calcium ATPases as described^13^. To expand the repertoire available for screening inhibitors and mutated sequences, several modifications were introduced in expression studies.

To compare with mammalian SERCA1a pump a new strain (K667::SERCA1a) was generated by transformation of K667 with plasmid pUGpd-SERCA1a. A vector-only control (K667::pUGpd) was included in experiments assessing selectivity and specificity of inhibitors. SERCA1a expression in yeast (Figure 1a) restores calcium tolerance (Figure 1b) to similar levels to that seen in Pulcini *et al*. for PfATP6 (Figure 2;^13^). Sensitivity to inhibitors is more apparent when they are used to inhibit yeast growth in the presence of calcium, at concentrations approaching those maximally tolerated by a particular strain. 20-40 mM total extracellular calcium concentration is optimal for assessing inhibition of PfATP6 (Figure 3;^13^). For SERCA1a any concentration greater than 50 mM (up to 250 mM) is sufficient to monitor complete inhibition (S. Figure 1b).

**Figure 1.**
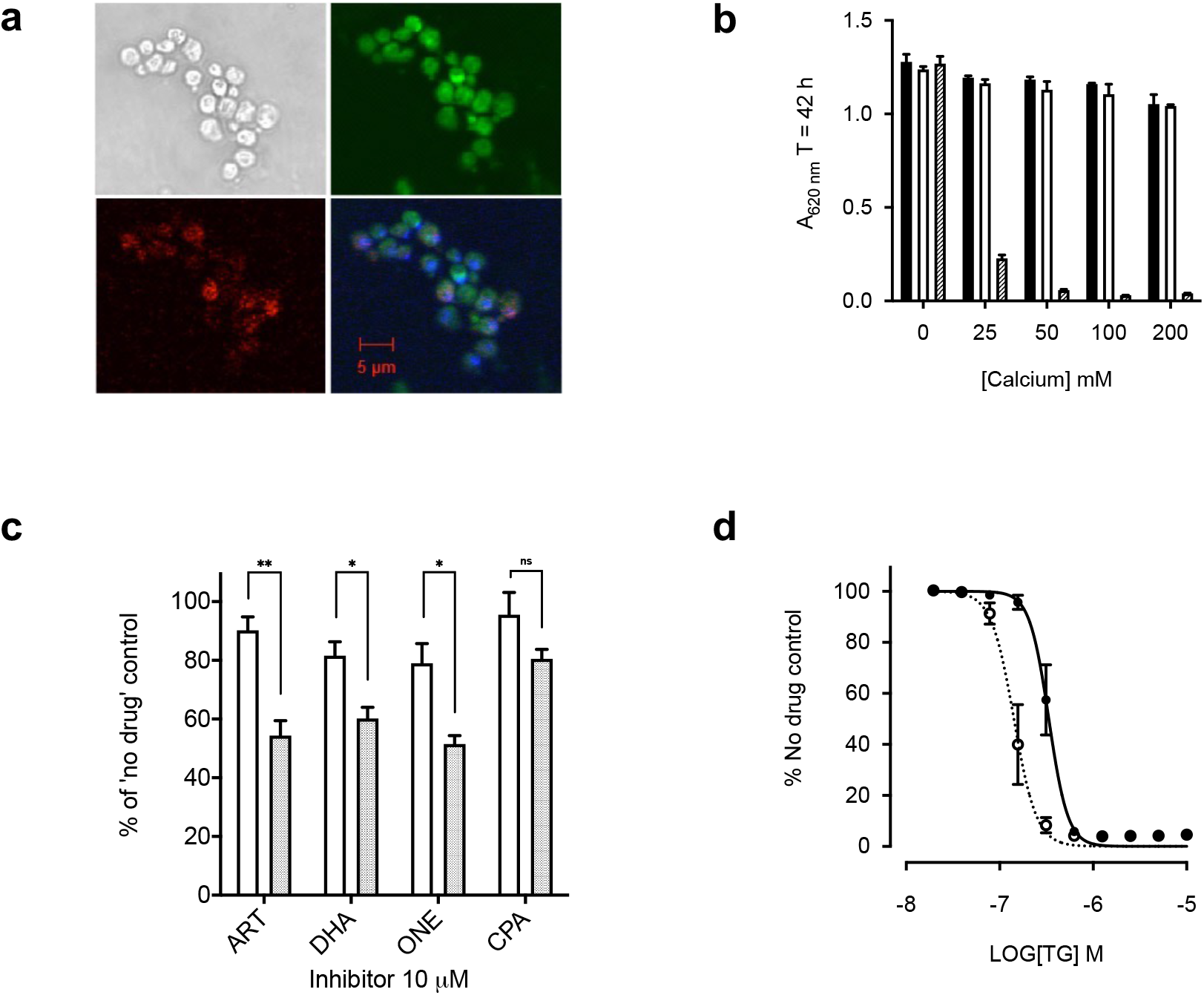
Yeast expression and optimisation. **A.** Immunofluorescence of SERCA1a. Yeast (top left) stained with primary SERCA1a antibody and Texas Red-tagged goat anti-mouse secondary antibody (bottom-left), as well as Di_6_OC stain for ER (top right), and DAPI (bottom right merge). **B.** Growth of K667[SERCA1a] strain (white), reference strain BY4741 (black), and K667[SERCA1a] treated with 1 μM thapsigargin (TG) (striped) in an extracellular calcium concentration range. Error bars are S.D. of the means of 5 technical replicates. **C.** K667[SERCA1a] strain (white) and K667[PfATP6] strain (shaded) treated with10 μM of artemisinin (ART), dihydroartemisinin (DHA), artemisone (ONE), and cyclopiazonic acid (CPA). Error bars are S.E.M. of 3 biological means of 5 technical replicates. (** = p < 0.0075, * = p < 0.05) **D.** Dose response curves of K667[SERCA1a] strain (solid line black square) and K667Δpdr5[SERCA1a] (dotted line open circles) to TG.

We confirmed that rescue in individual yeast strains was dependent on PfATP6 or SERCA1a by using cyclopiazonic acid (CPA) as a relatively unspecific inhibitor of both pumps. Artemisinin derivatives selectively inhibit PfATP6 function leaving SERCA1a unaffected at comparable concentrations (Figure 1c) and are used to establish specificity.

Yeast expresses ABC transporters that may modulate sensitivity to inhibitors in whole cell screening assays by enhancing their efflux. We knocked out *PDR5*, a key efflux pump, to establish its contribution to sensitivity of whole cell assays to inhibitors^20^. For *K667ΔPDR5::SERCA1a* the IC_50_ was over two-fold lower for thapsigargin (141.8 ± 43.9 nM) compared with IC_50_ in *K667::SERCA1a* (334.7 ± 67.5 nM; p = 0.015)(Figure 1d). The IC_50_ values for different artemisinin derivatives in this strain of yeast (*K667ΔPDR5::PfATP6*) were 4-11 times lower than those for the PDR5 intact strain *K667::PfATP6* (Table 1). *K667ΔPDR5* was therefore used for subsequent experiments.

**Table 1.** IC_50_ values for each artemisinin derivative for K667[PfATP6] and K667*Δpdr5*[PfATP6]. Significant difference between wild type and *pdr5* knockout for each derivative is indicated by asterisks (* p < 0.05;** p < 0.005; *** p< 0.0005). Means are of three biological replicates.

| Drug | K667[PfATP6] IC<sub>50</sub><br>(95% CI) | K667Δ*pdr5*[PfATP6]<br>IC<sub>50</sub> (95% CI) | Fold<br>difference |
| --- | --- | --- | --- |
| Artemisinin | 10.6 (8.2 – 13.8) | 1.8 (1.3 - 2.6) | 6** |
| Artesunate | 14.8 (10.0 – 21.9) | 3.7 (2.9 - 4.7) | 4* |
| Dihydroartemisinin | 23.8 (16.9 – 33.7) | 2.1 (1.6 - 2.8) | 11** |
| Artemether | 15.7 (12.5 – 19.7) | 3.4 (2.8 - 4.1) | 5*** |
| Artemisone | 3.6 (2.4 – 5.3) | 0.9 (0.6 - 1.3) | 4** |

### Mechanism of action investigations

Artemisinin’s mode of action is not yet fully elucidated, but it alkylates multiple proteins after being activated by haem in malarial parasites and in cancer cell lines^21^. To confirm PfATP6 is alkylated by artemisinins in yeast as well as parasites, we used a biotin-derivatised dihydroartemisinin (NewChem Technologies, Figure 2a) that retains antiparasitic potency (IC_50_ 5.0 nM ± 2.0 nM compared with dihydroartemisinin 2.5 nM ± 1.4 nM; p=0.15) to label PfATP6 or SERCA1a.

**Figure 2.**
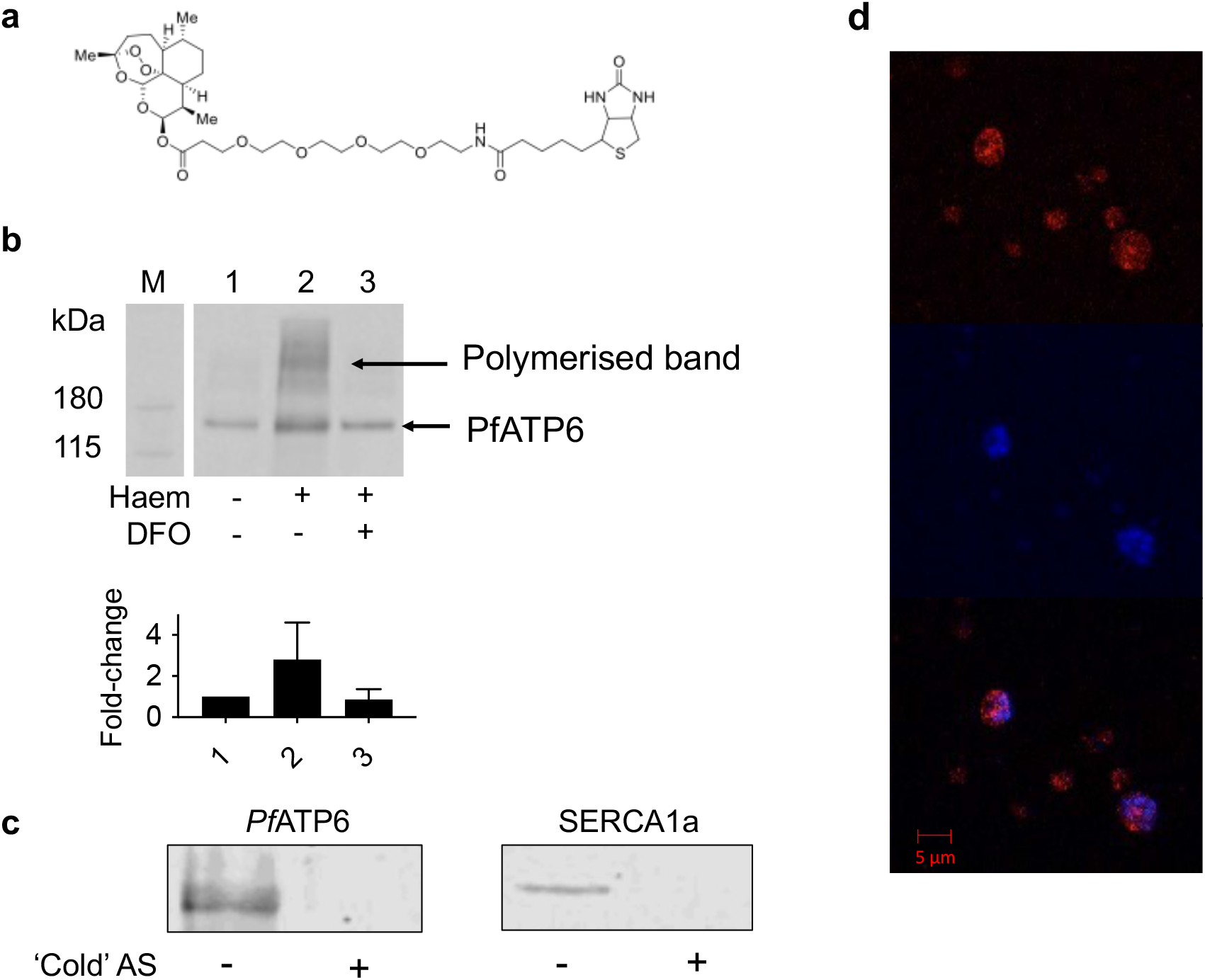
Artemisinins’ interaction with PfATP6. **A.** DHA-biotin probe structure. **B.** Western blots of PfATP6-enriched microsome pull-downs untreated (1), preincubated with 200 μM haem (2), and pre-incubated with haem and 200 μM DFO (3). Fold change refers to band intensity relative to PfATP6 (1). **C.** Western blots of PfATP6- and SERCA1a-enriched microsome pull-downs pre-incubated with ‘cold’ artesunate before performing pull-downs with DHA-biotin. **D.** Immunofluorescence assay with *P. falciparum* parasites. Trophozoite-stage parasites were pre-incubated with DHA-biotin before staining with TRITC-tagged anti-biotin antibody (red, top panel), and by DAPI (blue, middle panel). Merge in bottom panel.

PfATP6 binding to DHA was confirmed through pull-down and western blot analyses (Figure 2b), and mass spectrometry identified several proteins alkylated by the tagged dihydroartemisinin when applied to membrane extracts and whole yeast, including PfATP6 in K667[PfATP6] preparations. Yeast lacking PfATP6 were not labelled at the masses shown in Figure 2b (Table 2). SERCA1a could not be labelled to the same extent in parallel experiments (Figure 2c), consistent with a decreased potency of artemisinins against the native mammalian orthologue (Figure 1c). Pre-incubation of PfATP6 with excess dihydroartemisinin competed with the tagged dihydroartemisinin, preventing it from binding (Figure 2c). The addition of haem increased labelling of PfATP6 exposed to tagged dihydroartemisinin, as it does in parasites (Figure 2b and^18^). Conversely, chelation of Fe^3+^ using desferrioxamine (DFO) attenuates the interactions between artemisinins and targets (Figure 2b). Immunofluorescence assays with parasites using a FITC-labelled anti-biotin antibody localised the tagged dihydroartemisinin to the area surrounding the nucleus, most likely the ER, which is where PfATP6 is localised (Figure 2d)^13^.

**Table 2:** Mass spectrometry results of all proteins pulled down by DHA-biotin from both PfATP6-expressing whole yeast and PfATP6 enriched microsomes. List is separated into unassigned proteins, mitochondrial proteins, those also pulled down from *P. falciparum* and human cancer cells, and those also identified in the yeast microsomes.

| Unassigned proteins | Mitochondrial | Also identified in *P. falciparum*<br>(Wang, Ismail and Zhou) | Also identified in yeast protein-enriched<br>vesicles |
| --- | --- | --- | --- |
| Adh1p alcohol dehydrogenase | 6-phosphogluconate dehydrogenase | 40S ribosomal protein S21 | ABP PfATP6 |
| Ahp1p | Acc1p Acetyl-CoA carboxylase | 40S ribosomal protein S81 | Arc1p tRNA delivery |
| Arc1p tRNA delivery | Acetyl-coenzyme A synthetase | ABP PfATP6 | Elongation factor 1-alpha |
| Bgl2p endo-beta-1,3-glucanase | ATP synthase subunit alpha | Act1p actin | Hsp30p |
| Bmh1p | Cyc1p po-cytochrome c | ATP-dependent 6-phosphofructokinase | Plasma membrane ATPase |
| Bub2p | Cytochrome c oxidase subunit 2 (Fragment) | Elongation factor 1-alpha | DNA-directed RNA polymerase I subunit RPA43 |
| Dur1,2p urea carboxylase/allophanate hydrolase | Glutamate decarboxylase | Eno2p | Rps11bp |
| Erg20p synthesizes the Rer2p substrate farnesyl diphosphate | Glyceraldehyde-3-phosphate dehydrogenase | Eukaryotic translation initiation factor 5A | Sec28p coatomer subunit epsilon |
| Gag | Malate dehydrogenase | Hsp104p | Yro2p |
| Nam9p | Pck1p phosphoenolpyruvate carboxykinase PCK1 | Hsp12p |  |
| Pdc1p Peroxisomal protein of unknown function | Pet9p ADP/ATP transporter | Hsp26p |  |
| Rnr2p ribonucleotide-diphosphate reductase subunit RNR2 | Phosphoglycerate kinase | Hsp30p |  |
| Rpl10p ribosomal, chaperone | Por1p Mitochondrial outer membrane protein of unknown function | Ribosomal protein L19 |  |
| Rpl11ap | Pyruvate carboxylase | Rps18ap ribosomal subunit protein |  |
| Rpl16bp | Pyruvate kinase | Ssa1p ATPase HSP70 family |  |
| Rpl9ap | Qcr2p ubiquinol-cytochrome-c reductase subunit 2 |  |  |
| Rps14ap | Succinate dehydrogenase |  |  |
| Ubi4p ubiquitin binding | Superoxide dismutase [Cu-Zn] |  |  |
| Ugp1p UTP glucose-1-phosphate uridylyltransferase | Triosephosphate isomerase |  |  |
| Uncharacterized protein |  |  |  |
| Wtm1p Transcriptional modulator |  |  |  |

### Compound screening

Several individual compounds and three compound sets were screened using whole cell assays: the Medicines for Malaria Venture (MMV) malaria box library^22^, the MMV OZ box, and the thaperoxides^10^. The MMV malaria box is a set of 400 compounds with (sub)micromolar antimalarial activity.

Thaperoxides are derivatives of thapsigargin with an endoperoxide bridge introduced to resemble artemisinin more closely. The MMV OZ box is a blinded set of semi-synthetic artemisinin derivatives. To define assay robustness *Z*’ values were derived^23^, for PfATP6 inhibition with CPA *Z*’ = 0.97 ± 0.123, and for SERCA1a with TG *Z*’ = 1.00 ± 0.012.

Hits from the compound screen are summarised in Figure 3a. Initially six hits were identified from the MMV malaria box. Three of these hits gave results that were too variable to take forward. MMV665807 showed the highest reproducible potency against PfATP6 with almost no inhibition of control strains (*K667ΔPDR5::SERCA1a* and *K667ΔPDR5::pUGpd*). Hits from the malaria box were characterised in more detail (Table 3) in *PDR5*-knock-out strains and in cultured parasites. Identification of several new chemically unrelated classes of inhibitor of PfATP6 (Figure 3b and Table 3) confirmed a correlation between inhibitory constants for PfATP6 derived in yeast and anti-parasitic potency (Pearson’s r^2^ = 0.7 (n = 3; p = 0.004)).

**Figure 3.**
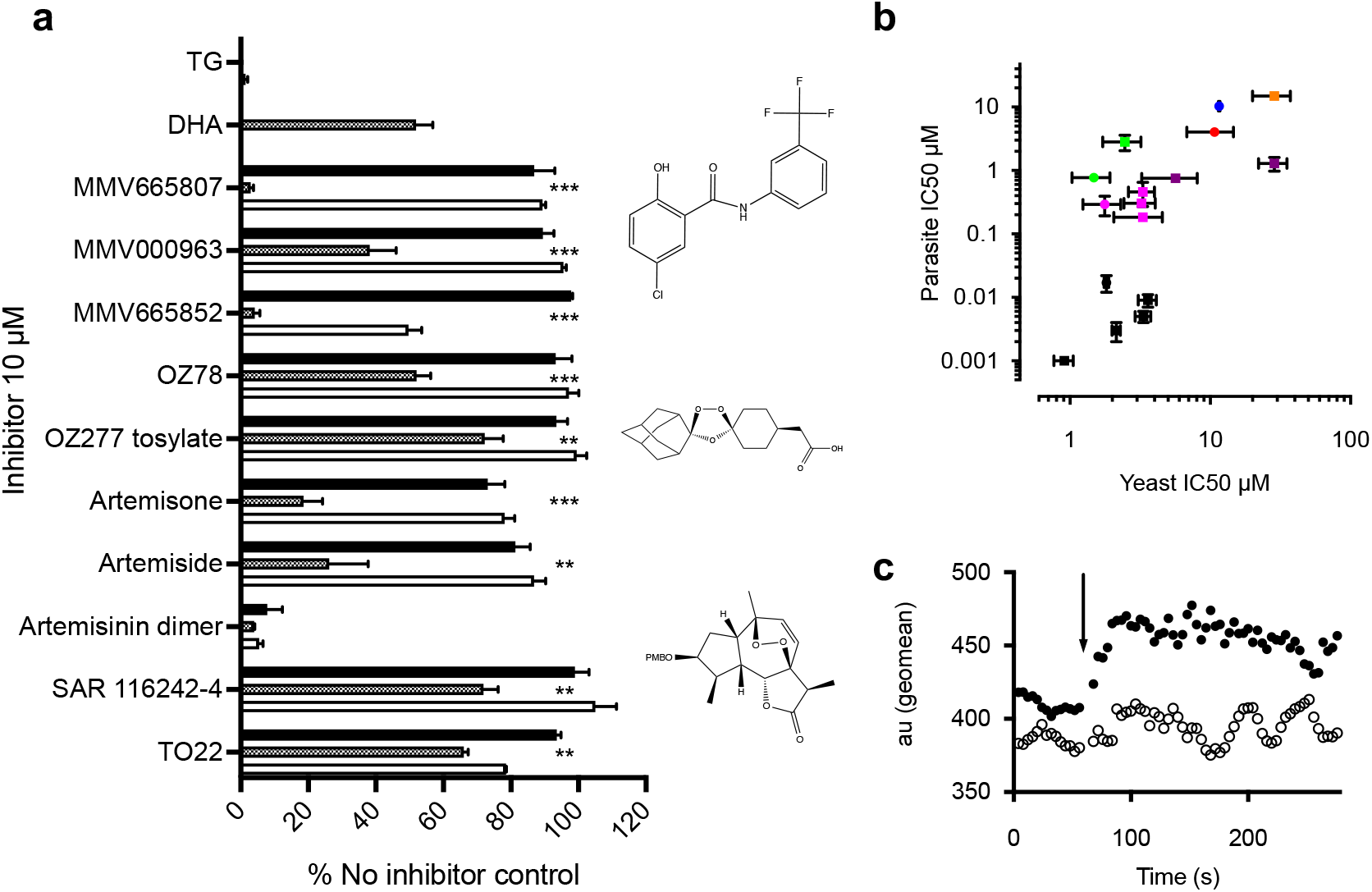
Inhibitor screens on whole yeast and parasites. **A.** All compounds were screened against K667Δpdr5[PfATP6] and two control strains K667Δpdr5[SERCA1a] and K667Δpdr5[pUG]. K667Δ*pdr5*[PfATP6] positive control is DHA and K667Δ*pdr5*[SERCA1a] is TG. For each class of inhibitor a representative structure is shown. **B.** The correlation between the yeast assay-derived IC_50_ values (μM) and the *in vitro* parasite assay-derived IC_50_ values (μM) for inhibitors had a r value of 0.7 (Pearson analysis P = 0.004). Artemisinins are in black. MMV compounds are in green, magenta, red/orange and purple. Circles denote parent compounds and squares derivatives. CPA is represented by the blue circle. Error bars are ± S.E.M. **C.** Parasites treated with artemisone had perturbed calcium transport. Arrow indicates addition of artemisone. Geometric means were derived from 3 biological replicates. P = 0.016 (one way ANOVA uncorrected Fisher’s LSD).

We next investigated if addition of artemisone (the most potent PfATP6 inhibitor and antimalarial) perturbed calcium homeostasis by measuring [Ca^2+^]_free_ in parasites using a cameleon-Nano biosensor^24^. Artemisone significantly increased [Ca^2+^]_free_ in parasites, confirming the relevance of inhibition of PfATP6 (Figure 3c).

Other compounds suggested by the literature were screened but had limited success: Hypericin, BHQ, saikosaponin, spiroindolones, and *tert*-butyl peroxide (Supplementary Figure 1a)^25–28^. Of these, saikosaponin and spiroindolones inhibited PfATP6 but only at high concentrations (>100 μM). These results are useful to train selection strategies for future compound screenings identifying PfATP6 inhibitors.

### Mutations on drug sensitivity

After expression in *Xenopus* oocytes, sensitivity of SERCA1a to artemisinins was increased by mutating a key amino acid residue (E255L) in the thapsigargin-binding pocket^29,30^. We confirmed that ^E255L^SERCA1a was more sensitive to all tested artemisinins, and conversely became five-fold less sensitive to thapsigargin *i.e*. IC_50_ = 793.6 ± 247.4 nM, versus 146.8 ± 43.9 nM for wild-type (p = 0.011) (Figure 4a and b respectively).

**Figure 4.**
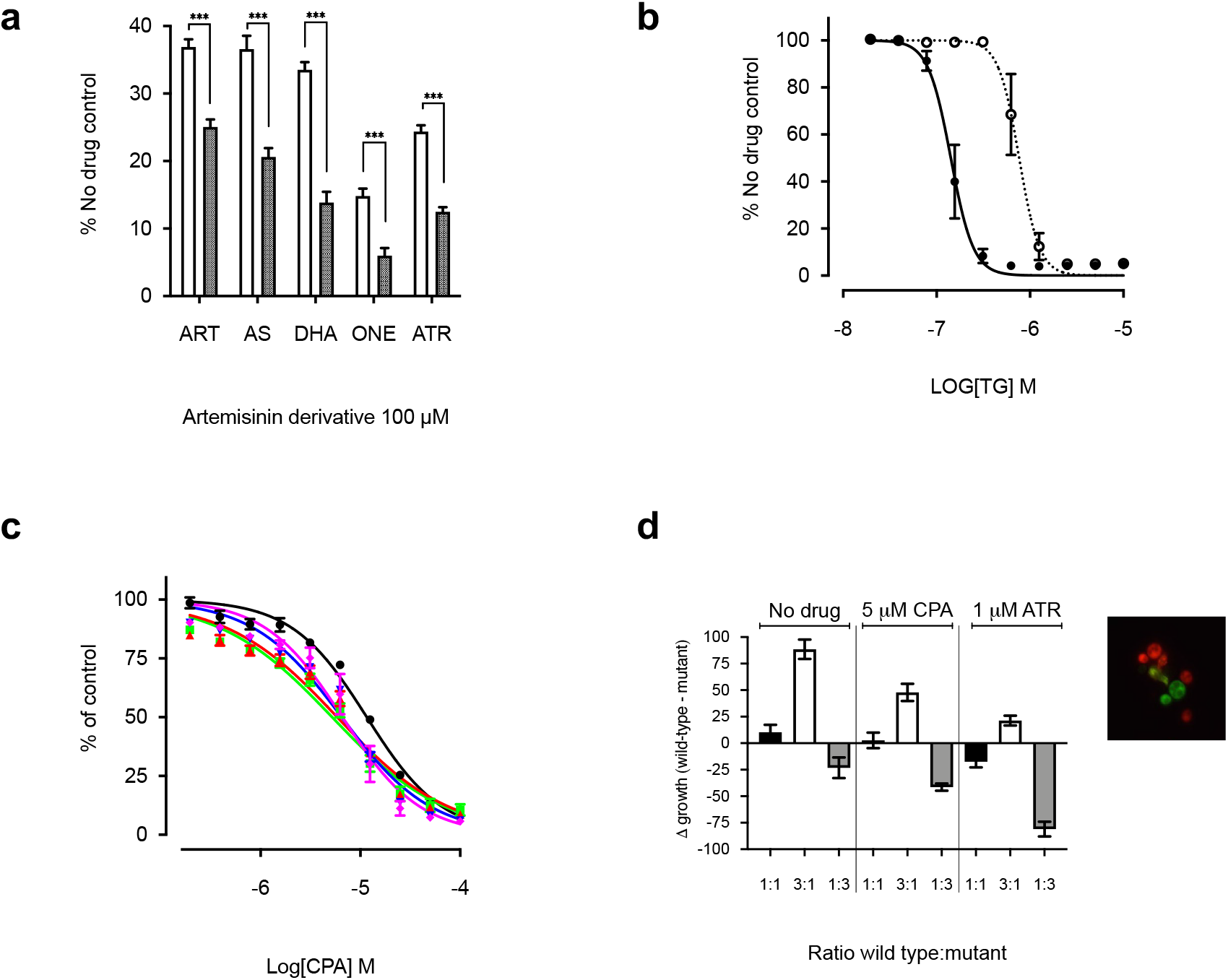
Effect of SNPs on artemisinin sensitivity. **A.** K667Δpdr5[SERCA1a]^E255^ (white) and K667Δpdr5[SERCA1a]^2551^ (black) were treated with 100 μM artemisinin (ART), artesunate (AS), dihydroartemisinin (DHA), artemisone (ONE) and artemether (ATR). *** = P < 0.0001 **B.** Dose response curve of K667Δpdr5[SERCA1a]^2551^ (dotted) and K667Δpdr5[SERCA1a]^E255^ (solid) to thapsigargin (TG). NB: IC_50_ for K667Δpdr5[SERCA1a]^E255^ same as that for Figure 1d, as the assays were performed together. **C.** Dose response curves to CPA for K667[PfATP6] wild type (black circles) versus strains expressing PfATP6 with resistanceconferring SNPs S769N (blue triangle), L263E (green square), A623E (red triangle), and A623E/S769N (magenta diamond). **D.** Fitness cost of K667Δpdr5[mCherry][PfATP6]^769N^ compared to the wild type K667Δpdr5[Venus][PfATP6]^S769^, with and without drug pressure from 1 μM artemether (ATR), or 5 μM CPA. Extracellular calcium concentration 50 mM. Fitness cost presented as change in growth between wild type and mutant after co-incubation. Differences in growth were calculated by the relative fluorescence of each strain when grown together at different starting ratios of wild type:mutant *i.e*. 1:1 (black), 3:1 (white) or 1:3 (grey).

### Fitness cost

Mutations in drug targets can have variable phenotypic manifestations in parasites due to opposing effects of decreased parasite fitness and decreased artemisinin sensitivity as previously proposed^31,32^. To test this hypothesis, we introduced *in vitro* artemether resistance-conferring mutations S769N and A623E in PfATP6 that have been previously identified in field isolates^31,33^. These naturally occurring mutations were also supplemented by an L263E mutation that we have previously shown to confer variable resistance in parasites to artemisinins, as it substitutes the malarial amino acid for the mammalian equivalent. Pulcini *et al* (Table 1;^13^) also showed that these mutations confer resistance to artemisinin and its derivatives in the yeast heterologous expression model. ^A623E^PfATP6 allowed the yeast to grow in all artemisinins at levels significantly higher than the wild type, except with artesunate. Similarly for ^S769N^PfATP6 with DHA. Otherwise, all mutations improved yeast growth in the presence of artemisinins.

Mutated PfATP6 sequences were also assayed in yeast for sensitivity to cyclopiazonic acid (CPA), which acts to inhibit SERCA pumps in a different way, both by interaction at different binding sites to those predicted for artemisinins and by hydrophobic interactions, not alkylation^34^. All mutations (L263E, A623E, S769N, A623E/S769N) increased sensitivity to CPA, both in parasites^13^ and the yeast heterologous expression assay (Figure 4c). Mean IC_50_s and S.D. were 11.59 μM ± 0.02 for wild-type, versus 5.21 μM ± 0.03 for L263, 5.86 μM ± 0.04 for A623E, 6.49 μM ± 0.03 for S769N, and 6.61 μM ± 0.041 for double mutant A623E/S769N. There was a significant difference between the wild type and all mutants (one-way ANOVA multiple comparisons, P value = <0.0001). These findings are consistent with the suggestion that mutations reduce the functionality of PfATP6 as L263E also does in parasites.

### Fitness in yeast

To examine the hypothesis that mutations in PfATP6 or SERCA1a decrease fitness in yeast rescue assays, we generated fluorescent reporter strains of yeast (Venus for wild-type PfATP6-expressing yeast and mCherry for the mutant PfATP6-expressing yeast; see image box in Figure 4d). These constructs were used in competition experiments between native and mutated PfATP6 sequences. As ^S769N^PfATP6 was less sensitive to artemether in parasites in French Guyana, this mutation was selected for detailed investigation^31^. Growth was unaffected by fluorescent markers compared to the non-fluorescent congenic strain (S. Figure 1b), confirming that markers do not cause growth disadvantage.

Across a range of extracellular calcium concentrations (optimal concentration 50 mM), native PfATP6 outcompeted ^S769N^PfATP6 in growth assays when experiments were begun at ratios of 1:1 or 3:1 (native:mutant strains) and grown until the yeast reached late log phase (18h). Growth levels were comparable when experiments began with 1:3 ratio of native to ^S769N^PfATP6. When selection with artemether (1μM: approximately yeast IC_80_) was applied under these conditions, the fitness cost of ^S769N^PfATP6 became attenuated, and yeast expressing this mutant had a growth advantage, especially when starting mixtures were in a ratio of 1:3 (native:mutant ratio; Figure 3d). When CPA was used (5 μM: approximately yeast IC_80_) instead of artemether as a selective agent, ^S769N^PfATP6 lost this advantage.

*In silico* modelling shows the changes in the druggable pockets with different mutations (Figure 5). The X-ray crystal structure of the SERCA in E2 (E309A mutant) was used as a template for building the homology models of PfATP6 and its mutations. The crystal structure shares 48% of sequence identity with pfATP6 and it covers amino acids from 5 through 1217. The predicted binding pocket for the modeled PfATP6, ^L263E^PfATP6, ^A623E^PfATP6, and ^S769N^PfATP6 has volume of 624.411, 922.372, 1746.135, and 1048.841 Å^3^ respectively. The mutations, even the distant ones, led to changes in volume of the binding cavity adjacent to Leu263.

**Figure 5.**
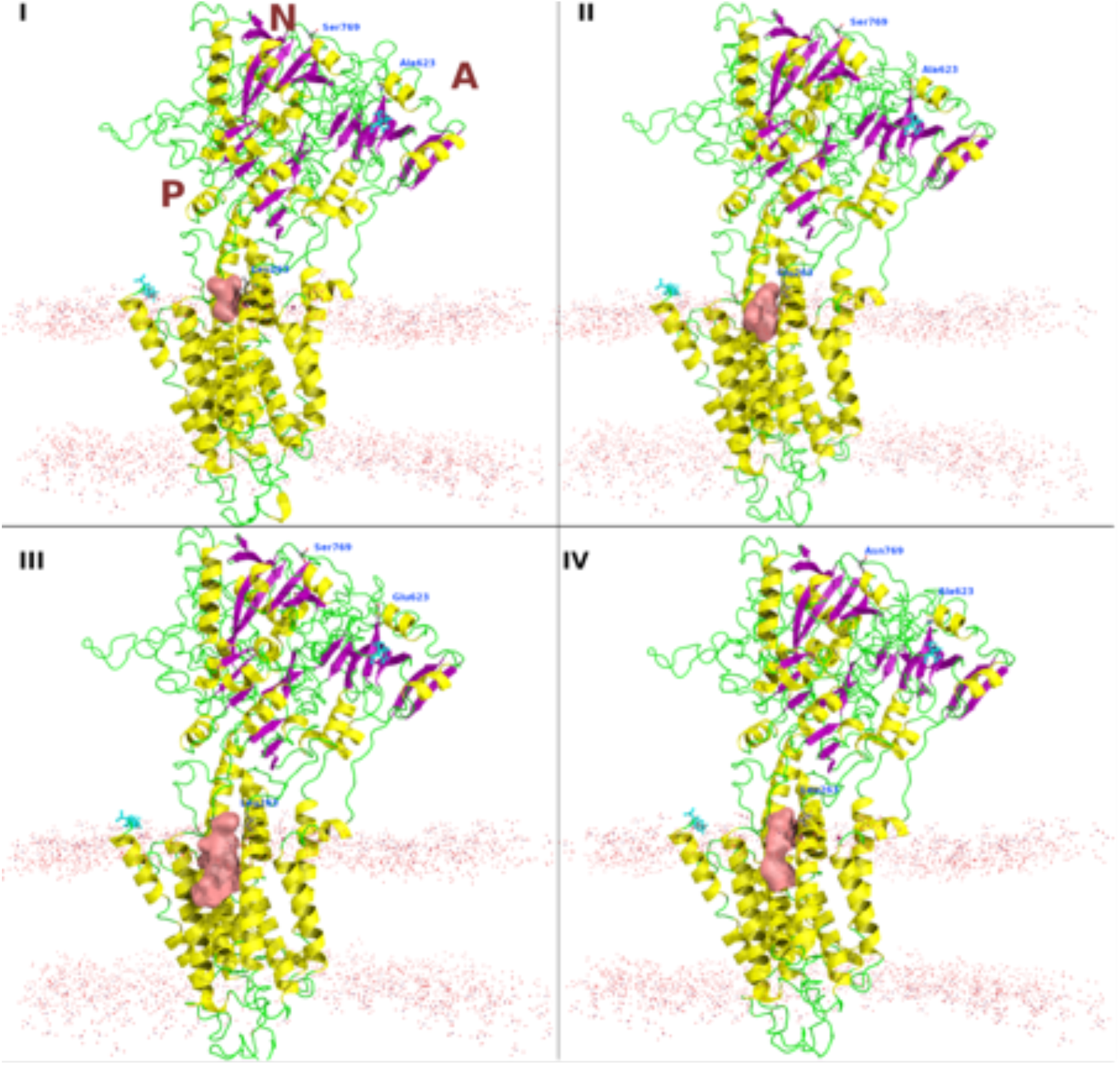
Homology models for the wild-type PfATP6 (I), ^L263E^PfATP6 (II), ^A623E^PfATP6 (III), and ^S769N^PfATP6 (IV). The druggable pocket close to Leu263 is shown as reddish surface. The membrane position is shown as dots. The N and C terminals are shown as cyan sticks. The cytosolic domains are labeled as N, A, and P, indicating the nucleotide-binding domain, the actuator domain, and the phosphorylation site respectively.

## Discussion

More than three decades ago we suggested that parasite integral membrane transport proteins may be useful drug targets^35,36^. In the course of investigating various families of transport proteins we first isolated, functionally characterised in *Xenopus* oocytes, and genetically validated the parasite’s hexose transporter, PfHT, as a new drug target^37^. Co-crystallisation of PfHT with the selective inhibitor we characterised has confirmed that structure-based design of inhibitors against integral membrane proteins of *P. falciparum* is feasible^38^.

Our early work also pointed to a family of P-type cation ATPases as promising for further study as drug targets. PfATP4 was characterised as an atypical cation ATPase initially proposed to be a calcium ATPase^39^ and then was found to export sodium from the intraerythrocytic parasite^40^. These studies are particularly interesting because PfATP4 is now an established drug target for several novel classes of antimalarial, including the spiroindolones that have reached clinical studies in development^41^. PfATP4 also has mutations that can confer resistance to inhibitors and drugs^42^.

PfATP6 has also been confirmed as a validated drug target by genetic studies^13^. Artemisinins were first shown to inhibit PfATP6 after expression in *Xenopus* oocytes, and to interact with PfATP6 in parasites^9^. These observations led to the hypothesis that PfATP6 was the key target of artemisinins’ antiparasitic activity. Since then, artemisinins have displayed promiscuity in their targeting of parasite proteins by alkylating them^18,19^. This mechanism may explain several features of assays with artemisinins. For example, parasites display both higher variances and inconsistencies in IC_50_ responses to artemisinins when bearing mutations in PfATP6^11^, predicted to reduce their susceptibility to artemisinins. The presence of more than one important target for artemisinins’ antimalarial activity (64-123 artemisinin-labelled proteins were identified, although not all would be plausible targets) may explain some variability in IC_50_ results because ‘secondary’ or ‘tertiary’ targets may become more important if a ‘primary’ target becomes less susceptible. This suggestion is reassuring for the risk of developing artemisinin resistance, because multiple targets may maintain parasite sensitivity to this class of antimalarial. Consistent with this suggestion of multiple targets, there may even be more than one binding site within a single protein target.

Different artemisinin derivatives behave differently against mutant PfATP6 *in vitro*. Adding to this complexity in interpreting susceptibilities of parasites with mutations in PfATP6 is their fitness cost, suggested in cultured parasites for those bearing the ^S769N^PfATP6, and now demonstrated in yeast models^32^. It also explains why it may have proved difficult to establish parasites with some PfATP6 mutations in cultures *ex vivo* for longer term further study. Studying the effects of mutations in PfATP6 observed in field isolates, and in laboratory models of parasites and after heterologous expression are consistent with these mutations being able to exert remote influence on the properties of drug binding regions (Figure 5). Expression in yeast suggests that mutations in the cytosolic domains of PfATP6 (S769N and A623E, for example) can also exact fitness costs, and these costs can may be ameliorated under drug selection pressure with some artemisinin derivatives.

To address differences in results of assays of PfATP6 in *Xenopus* oocytes reported by different groups, we developed a robust yeast whole cell assay to study artemisinins, to compare results to mammalian SERCA and to allow screening of inhibitors. Earlier immunolocalization experiments demonstrated PfATP6 is expressed in internal membrane structures in yeast^13^. The alkylation of PfATP6 by an artemisinin derivative when expressed in yeast (as in parasites), together with growth rescue of inhibition by artemisinins in this model confirms our earlier observations made in *Xenopus* oocytes on inhibition of PfATP6 and the effects of mutations^9,29^.

To confirm that mammalian SERCA1a can be made more susceptible to artemisinins, we also mutated its sequence as before (^E255L^SERCA1a) and again demonstrated increased susceptibility to artemisinins^29^ (Fig 4a) Interestingly, this sequence is less susceptible to thapsigargin (Fig 4b). These observations confirm that artemisinins target PfATP6, and that this inhibition can result in increases in free intraparasitic calcium concentrations when challenged with a potent artemisinin (Fig 3c). These observations also highlight how methodologies used to assay function in different expression systems may give apparently inconsistent results. *In vitro* activity assays also demonstrate this point, as no compound tested showed any inhibition of purified PfATP6 despite apparent inhibition of purified SERCA1a by CPA in parallel experiments, suggesting PfATP6 may not function in a representative way in *in vitro* assays (S. Figure 1c and 1d).

Treatment failure with artemisinin combination therapy (ACT) emerged in the early 1990s and is currently localised to the Greater Mekong region^43,44^. Fortunately, appropriately selected ACTs continue to be effective in treating *P. falciparum* infections in this region when the partner drug is efficacious, and case numbers continue to fall^45^.

Using PfATP6 as a ‘model’ target allowed assessment of the importance of other targets, because even with mutations that are predicted to decrease sensitivity of PfATP6, assays in parasites bearing such mutations suggested more variable IC_50_ results, both in field and laboratory generated strains. Fitness cost in the yeast model is ameliorated when these PfATP6 mutant strains are exposed to artemisinin derivatives, perhaps explaining why some parasites strains have been unstable in the laboratory when grown without drug selection pressure^32^. Future fitness cost assays could be carried out with yeast that have fluorescent markers incorporated into chromosomal DNA, rather than in a vector, to provide a simpler ‘plug-and-play’ platform.

To address problems of drug resistance in parasites, we exploited the yeast screening model, as it is eukaryotic, easily genetically manipulated, and knockout libraries are freely available (*e.g*. through EUROSCARF, Germany). One limitation is that any proposed target needs to have an orthologue in yeast, so targeting mechanisms that relate to parasite-specific properties such as red cell invasion might be difficult. Also, anti-parasitic drugs are often stage-specific (which needs to be taken into account when choosing targets, since yeast do not have comparable life cycles). Yeast can respire aerobically or anaerobically, depending upon the carbon source in the medium, further distinguishing it from parasites and an inhibitory effect on yeast aerobic respiration apparati is only apparent when the carbon source ensures yeast rely on mitochondrial respiration, rather than glycolysis^46^.

Inhibitory constants in yeast screens were correlated with those in parasite assays. Differences in their magnitude (*e.g*. artemisinin = 17 nM in parasites vs. 1.8 μM in yeast) could be explained by several factors. A requirement for activation of artemisinins by haem is a possibility because haem was not included in the yeast growth assays. Haem abolishes artemisinins’ ability to inhibit yeast growth in fermentable media^46^. Also yeast has many efflux pumps that lower its sensitivity to inhibitors, and *K667ΔPDR5*::HIS[PfATP6] yeast was 4-11 times more sensitive to artemisinins than the K667[PfATP6] strain. We did not knock out other potential efflux pumps, of which more than a dozen remain. This mechanism and its relevance to parasites is amenable to further study by introduction, for example, of the *P. falciparum* efflux pump Pfmdr1.

In conclusion, several independent experimental findings are consistent with the hypothesis that artemisinins target PfATP6. PfATP6 can be inhibited by a variety of unrelated compounds after heterologous expression in a yeast screening system, demonstrating its potential as a drug target, and potencies correlate with parasite killing. Mutations in PfATP6 can incur fitness costs. Because artemisinins interact with multiple targets including PfATP6, our findings suggest that a decreased sensitivity to artemisinins (assayed in ring stages) may depend on a global response of the parasite by selection to delay its developmental cycle. They also suggest that the term ‘artemisinin resistance’ may be best reserved for parasites that have been selected to demonstrate increased IC_50_ values to artemisinins in conventional *in vitro* assays, rather than those focused on decreased ring stage sensitivity.

## Materials and Methods

### Reagents

All reagents were purchased from Sigma-Aldrich except for thaperoxides (made by M. Avery). The OZ derivatives and the malaria box compounds were kindly donated by the Medicines for Malaria Venture^22^. Reference strain and K667 strain were from EUROSCARF (Germany) and the plasmid containing SERCA1a was kindly donated by Prof. Ghislain (Université Catholique de Louvain, Louvain-la-Neuve, Belgium). Plasmids with fluorescent markers were kindly donated by Dr. Elizabeth Bilslan^20^. All kits were purchased from Qiagen. All mass spectrometry was performed by Dr. Jigang Wang. Antibodies were purchased from Abcam.

### Yeast strains

| Name | Strain | Genotype |
| --- | --- | --- |
| Reference strain | BY4741 | Mat a; *his3Δ1*; *leu2Δ0*; *met15Δ0*; *ura3Δ0* |
| K667 | W-303 | Mat a; *ade2Δ1*, *can1Δ100*, *his3Δ11*, 15 *leu2Δ3*, 11, 2, *trp1Δ1*, *ura3Δ1*, *cnb1::LEU1* *pmc1::TRP1* *vcx1Δ* |

### Transformations

Yeast transformations were performed using the method described by Gietz and Schiestl^47^ and transformants were selected by an uracil auxotrophic marker. Transformations were confirmed by colony PCR. Both K667 and *K667Δpdr5::HIS* were transformed with the following plasmids: pUGpd plasmid as a vector-only control (derived from pRS316^48^); br434 plasmid (derived from pRS316,^49^) containing the SERCA1a coding region from rabbit skeletal fast-twitch muscle (*pbr434-SERCA1a*); and pUGpd containing the yeast-optimised coding region of PfATP6 from *Plasmodium falciparum* (*pUGpd-PfATP6*). Site-directed mutagenesis was performed on the SERCA1a coding region to produce the mutation E255L in the protein. Mutations were also introduced into the PfATP6 coding region to produce mutations L263E, A623E, S769N, and the double mutation A623E/S769N^13^.

### Homologous recombination

To knock out the *PDR5* gene from the K667 strain of yeast, the auxotrophic marker HISMX was amplified using the following primers^20^: 5’-AGACCCTTTTAAGTTTTCGTATCCGCTCGTTCGAAAGACTTTAGA<u>ATGGCAGAACCAGCC</u>-3’ 5’-TGTTTATTAAAAAAGTCCATCTTGGTAAGTTTCTTTTCTTAACCA<u>TACTTCACATCAAAA</u>-3’ where the underlined region is homologous with HISMX. PCR products were used to transform the K667 strain as above. Knockouts were selected by their ability to grow on histidine-depleted selective medium. Recombination was confirmed with colony PCR.

### Immunocytochemistry

Immunocytochemistry was performed as described in Kilmartin and Adams^50^. Briefly, yeast were grown to an optical density equivalent to log phase and were prepared for immunostaining by removing the cell wall. The cell suspension was spotted on polylysine-coated microscope slides and dried, fixed, and permeabilised. Primary antibody (mouse anti-SERCA1a, Abcam) was added and incubated overnight at +4 °C. Secondary antibody (goat anti-mouse Texas Red-tagged) was added and incubated for at least two hours. The slides were stained with 5 μM Di_6_OC for 20 minutes to stain endoplasmic reticulum. Antifade mounting medium plus DAPI was added before sealing the slides. Yeast were visualised with a Zeiss Confocal LSM 510 microscope. Immunofluorescence with *P. falciparum* parasites was performed as in Pulcini *et al*.^13^, using paraformaldehyde/glutaraldehyde fixation method. Parasites were labelled as in Wang *et al*.^18^, using 500 nM of DHA-biotin probe. Parasites were visualised with a Zeiss Confocal LSM 510 microscope.

### Site-directed mutagenesis

The E255L mutation was introduced into SERCA1a using the following primers: 5’-GCCGCTGCAGCAGAAGCTGGATTTATTCGGGGAGCAG-3’ Forward 5’-CTGCTCCCCGAATAAATCCAGCTTCTGCTGCAGCGGC-3’ Reverse Site-directed mutagenesis was performed using the Agilent Quikchange Lightning site-directed mutagenesis kit following manufacturer’s instructions. Mutants were screened by Sanger sequencing (Eurofins, UK).

### Whole-cell screening assay

The screening assay used here was developed from the protocol used in Pulcini *et al*.,^13^. A single colony of the yeast strain to be screened was picked and grown in 5 ml of selective medium until stationary phase was reached. The culture was diluted 100-fold in ampicillin-supplemented YPD medium to an O.D._620nm_ equivalent to that of lag phase. YPD was supplemented with an appropriate concentration of calcium, depending upon the transformed strain. The optimum calcium concentration for the PfATP6-expressing strain was 22 mM, and 100 mM for the SERCA1a-expressing strain. The optimum calcium concentration was periodically monitored and, if necessary, adjusted using a calcium concentration range of 10 to 30 mM. A final volume of 200 μl of the yeast culture was added to each well of a 96-well plate, with 5 technical replicates included per inhibitor. All inhibitors were dissolved in DMSO unless otherwise stated. Stock solutions of each inhibitor were diluted 100-fold in each well to give the desired final concentration. Where IC_50_s were being determined, inhibitors were two-fold titrated from stock solution. The 96-well plates were incubated at 30°C for 42 hours. The yeast growth was estimated from the absorbance at 620 nm in a Tecan plate reader. Growth was normalised with the no-drug control to present data as % growth.

The data collected were analysed using GraphPad Prism 9.0, (GraphPad Software, CA, U.S.A).

### *In vitro* parasite assays

Inhibitor IC_50_s were calculated in parasite assays following the protocol described in Desjardins *et al*.^51^. Briefly, cultures were synchronised to ring stage by sorbitol lysis, using 5% sorbitol solution. The parasites were diluted to a final parasitaemia of 1%. Hypoxanthine-free medium was added to give a haematocrit of 4%. A two-fold dilution series of the inhibitors in hypoxanthine-free medium was prepared. 100 μl of each inhibitor concentration was seeded in a 96-well plate, with five technical replicates of each included. Two no-inhibitor controls *i.e*. hypoxanthine-free medium only, were included. Plates were incubated at 37°C in 5% CO_2_ for 24 h. After 24 h [^3^H]hypoxanthine was added to each well to a final concentration of 0.5 μCi/well and incubated for a further 24 h. Plates were then freeze-thawed to lyse the cells, and the [^3^H]hypoxanthine uptake was measured in a Beckman scintillation counter. Parasite calcium homeostasis was measured as described in Pandey *et al*.^24^, using artemisinin derivative artemisone.

### Fitness cost assays

Fitness cost assays were performed in yeast by transforming the strain K667[PfATP6]^wt^ with yEpGAP-Venus, and the strain K667[PfATP6]^769N^ were transformed with yEpGAP-Cherry. Both vectors were kindly donated by Dr Bilsland^20^. Growth of the transformed and untransformed strains were compared using growth curves, performed as in Moore *et al*.^46^. Standard curves were generated for each strain by measuring fluorescence at each optical density in a two-fold dilution series of the yeast. The yeast were diluted to an optical density equivalent to log phase and co-inoculated at an equal starting optical density. Yeast were incubated at 30°C for 24 hours at which point the yeast were in late log phase. The fluorescence was then measured for both fluorophores and growth was estimated by calculating the optical density using the standard curve. The optical density was also measured. The experiments were repeated in the presence of drug pressure using 1 μM Artemether or 5 μM CPA (*i.e*. approximate IC_10_).

Standard curves calculated from optical density measurements and fluorescent measurements of titrations of each fluorescent strain of yeast were used to relate relative fluorescence to the equivalent optical density, such that each strain in a 1:1 mixture should have equivalent relative fluorescence and make up 50% each of the total optical density.

### *In silico* modelling

Prime was used to construct and refine the 3D models of ^wild-type^PfATP6, ^L263E^PfATP6, ^A623E^PfATP6, and ^S769N^PfATP6 (Template: PDB accession code 5ZMV). The basic local alignment search tool of homology search was used to identify the highest homologous protein structures from the PDB repository (http://www.rcsb.org) by using position-specific iterative basic local alignment search tool, NCBI NR database, BLOSUM62 similarity matrix, gap opening cost of 11, gap extension penalty of 1, inclusion threshold of 0.005 and three iterations. The secondary structure prediction was then established by SSPro, followed by sequence alignment with ClustalW. Knowledge-based 3D model builder was used to construct the models for each target sequence. Loops were refined by using VSGB solvation model and extended serial loop sampling. Side chains were then re-predicted for the refined loops, and the final 3D models was energy minimized using OPLSe force field and VSGB solvation model. The protein structure quality was checked in the Schrodinger suite. The binding pockets of PfATP6 and all mutations were predicted using f-pocket to identify the most plausible cavities close to L/E263.

### Microsome preparation

Protein-enriched microsomes were prepared as described in Nakanishi *et al*.^52^, and Hwang *et al*.^53^. Briefly, yeast was grown in 1L of YPD medium until an O.D_.600nm_ of 1.5-2.0 was reached. Yeast were pelleted and resuspended in 50 ml per 500ml culture of buffer 1 (0.1M Tris-HCl, pH 9.4, 50 mM 2-mercaptoethanol, 0.1 M glucose). After shaking at 30°C for 10 min, the yeast were pelleted and resuspended in 50 ml per 500 ml original culture of buffer 2 (0.9 M sorbitol, 0.1M glucose, 50 mM Tris-Mes, pH 7.6, 5 mM DTT, 0.5x SD medium, 0.05% w/v Zymolase 20T) and incubated at 30°C for 1-2 h, with gentle shaking. The yeast were pelleted and washed with 30 ml per original culture of 1 M sorbitol. The yeast were pelleted and resuspended in 20 ml per original culture of buffer 3 (50 mM Tris-Mes, pH 7.6, 1.1 M glycerol, 1.5% w/v PVP 40,000, 5 mM EGTA, 1 mM DTT, 0.2% w/v BSA, 1 mM PMSF, 1x protease inhibitor) on ice and homogenised. The homogenate was pelleted and the supernatant was transferred to ultracentrifuge tubes. The pellet was resuspended in buffer 3 and pelleted again. This second supernatant was pooled with the previous and centrifuged at 150,000 *g* for 45 min at 4°C. The pellet was resuspended in 1-2 ml buffer 5 (5 mM Tris-Mes, pH 7.6, 0.3 M sorbitol, 1 mM DTT, 1 mM PMSF, and 1x protease inhibitor) and aliquoted for single use before freezing in liquid N_2_.

### Mass spectrometry and activity assays

Both whole yeast and microsomes were prepared for mass spectrometry analysis by adding 500 nM of probe to 200 uL (~0.6 mg protein) of yeast microsomes preparation, or 10 μM probe to 50 – 100 ml yeast culture at O.D. 1.0 - 1.5 (~35 - 70 mg total protein), and incubated for 4 hr at 30°C. For competition studies, x25 artesunate was added and incubated for 30 minutes, before adding the probe and incubating for 4 hr, at 30°C. Samples were then acetone-precipitated by adding 4 volumes of −20°C acetone:water (4:1) to the yeast pellet or microsomal pellet and incubating for 2 hours to overnight at - 20°C. Then the samples were pelleted at 13,000 – 16,000 x *g* for 10 minutes at 0 - 4°C and washed twice with −20°C acetone:water solution. After a final centrifugation pellets were dried in a freeze-drier for one hour. Subsequent pulldowns and LC-MS/MS was performed as described in Wang *et al*.^18^.

Activity assays were performed as in Longland *et al*.^54^ and LeBel *et al*.^55^, except the reactions were incubated at 30°C instead of 37°C. Briefly, microsomes were diluted to 75 μg/ml, with free calcium at approximately 1 μM, in 2 ml assay buffer (45 mM HEPES/KOH pH 7.2, 6 mM MgCl_2_, 2 mM NaN_3_, 250 mM sucrose, 12.5 μg/ml A23187, 2 mM EGTA, ± 2mM CaCl_2_) and incubated for 10 minutes at 30°C in presence/absence of inhibitor (10-100 μM). Then 5.8 mM ATP was added and incubated for a further 15 minutes at 30°C. The reaction is stopped with 0.5 ml 6.5% TCA, on ice, and centrifuged at 4°C for 10 minutes (16,000 xg). 0.5 ml of the supernatant was added to 1.5 ml of copper acetate buffer (0.25% copper sulphate pentahydrate, 4.6% sodium acetate trihydrate, dissolved in 2M acetic acid pH 4.0) and mixed by vortexing, before addition of 0.25 ml of 5% ammonium molybdate, followed by 0.25 ml METOL buffer (2% p-methyl-aminophenol sulphate, 5% sodium sulphite). Samples were incubated for 10 minutes and absorbance measured at 870 nm in spectrophotometer. Pi standard was 0.4387 g/100 ml KH_2_PO_4_.

### Pull-down assays

A 0.3 mg aliquot of PfATP6-enriched membrane vesicles, as well as an aliquot of SERCA1a and vector-only membrane vesicles, were thawed on ice. Half of each aliquot (0.15 mg) was preincubated with 12.5 μM artesunate at 30°C for 30 minutes. Then all samples had 500 nM DHA-biotin probe added (NewChem Technologies ltd) and incubated at 30°C for 4 hours. The samples were then pulled down using Dynabeads following manufacturers’ instructions. 2 mg of beads were used per sample. Beads and samples were boiled for 5 minutes in 0.1% SDS to separate the beads from the proteins and supernatants were run on a polyacrylamide gel. Western blots were performed according to the Thermo Fisher NuPage western blot protocol, using 10% Bis-Tris gels (Thermo Fisher, UK) with mouse anti-SERCA1a primary antibody or goat anti-PfATP6 antibody (1/1000), and LiCor donkey anti-mouse or donkey anti-goat fluorescent-tagged secondary antibodies (1/10,000). Blots were analysed using the Odyssey scanner system. Densitometry was performed using Image Studio software.

### Statistics

T-tests and one-way ANOVA were performed using Graphpad Prism software, unpaired and two-tailed. * = p <0.05, ** = p<0.005, *** = p<0.0005 except where stated otherwise.

## Acknowledgements

We gratefully acknowledge support from the European Community’s Seventh Framework Programme (FP7/2007-2013); NanoMal under grant agreement number 304948 to S. Krishna and H. Staines. H. Staines is supported by the Wellcome Trust Institutional Strategic Support Fund (204809/Z/16/Z) awarded to St. George’s University of London. S. Krishna is co-chair of the Guidelines Development Group for Antimalarials of the World Health Organisation. Views here are personal and do not represent those of the Committee. The authors have no other relevant affiliations or financial involvement with any organization or entity with a financial interest in or financial conflict with the subject matter or materials discussed in the manuscript apart from those disclosed.

## Abbreviations

PfATP6: *Plasmodium falciparum* ATPase 6
ACT: Artemisinin Combination Therapy
SERCA1a: Sarco/Endoplasmic Reticulum Calcium ATPase 1a
DHA: Dihydroartemisinin
PMC1: Calcium ATPase in vacuole
PMR1: Calcium ATPase in golgi
CPA: cyclopiazonic acid
DFO: Desferrioxamine
MMV: Medicines for Malaria Venture
BHQ: Benzohydroquinone
ART: Artemisinin
ATR: Artemether
ONE: Artemisone
AS: Artesunate
TG: Thapsigargin
PfHT: *Plasmodium falciparum* Hexose Transporter
PfATP4: *Plasmodium falciparum* ATPase 4
PfMDR1: *Plasmodium falciparum* MultiDrug Resistance transporter 1
DTT: Dithiothreitol
BSA: Bovine Serum Albumin
PMSF: Phenylmethylsulfonyl Fluoride
O.D.: Optical Density
TCA: Trichloroacetic Acid

